# A generative model for dimensionality reduction with millions of features and few samples

**DOI:** 10.64898/2026.08.04.742788

**Authors:** Corrado Pancotti, Piero Fariselli, Jonas Meisner, Anders Krogh

**Affiliations:** Helmholtz Munich, Helmholtz AI, Ingolstadter Landstraße 1, Munich, 85764, Germany; Department of Medical Sciences, University of Torino, Torino, Italy; Novo Nordisk Foundation Center for Basic Metabolic Research, University of Copenhagen, Copenhagen, Denmark; Department of Computer Science, University of Copenhagen, Copenhagen, Denmark; Center for Health Data Science, University of Copenhagen, Copenhagen, Denmark

## Abstract

**Motivation:** In this paper, we demonstrate that it is feasible to train a deep generative model for dimensionality reduction with millions of features using few samples, which makes this type of generative model a more versatile alternative to standard methods for dimensionality reduction. Specifically, we hypothesize that for a decoder-only model, the number of training samples required is almost independent of the feature dimensionality in most network architectures.

**Results:** Through an extensive set of experiments on synthetic non-linear data, we validate this hypothesis. We also train the model on a downsampled version of the 1000 Genomes Project (1KGP) dataset to further assess its behavior under controlled reductions in sample size. Furthermore, we train a deep generative decoder (DGD) on a curated dataset from the International Cancer Genome Consortium (ICGC), which contains 4.4 million features. It is trained on approximately 4,000 samples and tested on 1,000 samples. The resulting latent representation exhibits clear clustering, and when methods are reduced to the same number of dimensions, it outperforms PCA and VAE for tumor type classification. Additionally, the DGD is computationally efficient and can be trained on a 16GB GPU.

**Availability and implementation:** Code is available at https://github.com/cpancott/ReceptiveDGD.

**Supplementary information:** Supplementary data are available with this preprint.

## 1 Introduction

Many dimensionality reduction methods exist. The most common is principal components analysis (PCA) [1], which linearly projects data points onto a lower-dimensional space in a way that best preserves variation in the data. Common non-linear techniques include UMAP [2] and t-distributed stochastic neighbor embedding (t-SNE) [3], which both map onto a low-dimensional space in a way that attempts to preserve the local structure of the data. The non-linear methods are slow or impossible to use directly on large data sets of very high dimensions (thousands or millions of dimensions), and it is common to first do PCA to a few hundred dimensions and then apply the method on the PCA projection.

Another approach is to use an auto-encoder neural network [4, 5], or the generative variant called the Variational Autoencoder (VAE) [6]. These methods yield a low-dimensional representation in a hidden (latent) layer. One advantage of these methods compared to other methods is that one can design the optimal decoder (and encoder) to match the data. For instance, one would use a convolutional network for image data, a Bernoulli distribution for the output for binary data, and a Poisson distribution for the output for count data. This is in contrast to PCA, which is optimal for real-valued data with Gaussian noise and often gives bad results on, e.g., sparse binary data. Additionally, the autoencoder yields an easy way to map to the representation space and back to feature space.

An important question is how many samples are needed to make a meaningful dimensionality reduction. An *m*-dimensional PCA can be calculated from *m* + 1 samples regardless of the feature dimension (*d*), although more points are needed, of course, to obtain a meaningful result for noisy data. Similarly, the non-linear methods also work with relatively few samples and are not so sensitive to the feature dimension. In autoencoders, on the other hand, the number of parameters scale with the feature dimension, and therefore may seem to require more data.

Based on a simple theoretical approach, we argue from reasonable calculations that the number of samples needed to train a decoder-only generative neural network is essentially independent of the feature dimension for networks with the number of parameters proportional to the feature dimension. We show that it holds for simulated data in increasingly large dimensions and for data from the 1000 Genomes Project with up to almost 700 thousand features. Finally, we show results of a dimensionality reduction for a set of somatic cancer mutations with more than four million features and about 4 thousand samples.

## 2 Background

The purpose of dimensionality reduction methods is to map data points ***x*** ∈*R*^*d*^ to points (representations) in a low-dimensional space ***z*** ∈ *R*^*m*^ (*m < d*), in a way that, loosely speaking, minimally disturbs their distribution. Here, the original space *R*^*d*^ is called the feature space, and the low-dimensional space the representation space (or latent space).

PCA can be obtained by diagonalizing the data covariance matrix. Similarly, many other methods for dimensionality reduction minimize some loss dependent on the pairwise distances between points in feature space and representation space. These methods include multidimensional scaling [7] and t-SNE [3]. Other methods like Isomap [8] and UMAP [2] also use pairwise distances combined with a neighborhood graph. Such methods are computationally intensive, because they are considering the *N* ^2^ pairs, and with the exception of PCA, they are not applicable for a large number of high dimensional samples.

The linear subspace corresponding to the *m* top principal components (the principal subspace) can be found by a linear neural network minimizing the squared loss [1, 9]. Similarly, the auto-encoder approach optimizes a loss that depends on single samples and not a pairwise measures and they therefore have a lower computational complexity than those based on pairwise distances.

### 2.1 On the number of samples needed

A model is theoretically unidentifiable if the parameters cannot be uniquely inferred from your dataset. With non-linear models it is usually not possible to asses exactly, and we instead use “practical unidentifiable”, which means that the model is highly sensitive to noise or small changes in data [10].

Assume you have a feed-forward neural network with *m* inputs and *d > m* outputs and a total number of parameters, *C*. To determine the number of samples needed to train such a network, previous work [9] has argued that one should compare the number of parameters *C* to the number of constraints imposed by the training data, which is *dN*, where *N* is the number of samples. In other words, one needs *dN > C* samples, so the number of samples should be larger than *N*_*c*_ = *C/d*. For a generative model [11] where the *m*-dimensional representations are learned by maximum likelihood at the same time as the neural network (decoder), there are an additional *mN* free parameters, in which case *dN*_*c*_ = *mN*_*c*_ + *C*, or

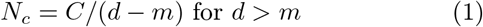

If we have a single hidden layer of size *h* in a fully connected decoder, then *C*≃ (*m*+*d*)*h*, so if *d* ≫ *m*, then *N*_*c*_ ≃ *h*, so the number of samples needed is independent of *d*. This holds in general for networks with a number of parameters proportional to *d*. If *C* = *cd*, where c is a constant, Equation 1 yields

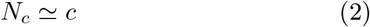

when assuming that *d* ≫ *m*. It shows that for most reasonable decoder architectures, we would expect that the number of samples needed is independent of the dimension of the feature space.

In this work we also use networks which are not fully connected, but with a “receptive field” of some size *w*. Let us say these receptive fields have a stride of *s*, then the number of units in the last hidden layer is *d/s* and the number of parameters in this layer is *wd/s*. Suppose the number of parameters of the rest of the neural network is *C*^*′*^, so the total is *C* = *C*^*′*^ + *wd/s*. From Equation (1) we then get *N*_*c*_ ≃ *w/s* + *C*^*′*^*/d* if we assume that *m* ≪ *d*. If *C*^*′*^ grows at most ∝ *d, N*_*c*_ is constant or even declines with increasing feature dimension.

These are not exact calculations, but more ballpark estimates. The main point is that the number of samples needed is expected to be essentially independent of the output dimension *d*.

For auto-encoders, the calculations are more tricky. If they are unconstrained, the encoder requires a much larger sample to be well specified [9], but the VAE is heavily regularized by the Kullback-Liebler term, which seems to prevent over-fitting, but then also sometimes increases the reconstruction error.

## 3 Simulations on synthetic data

### 3.1 Data generation

We generated non-linear synthetic data points *d*-dimensional space distributed around spherical clusters. In particular, these points are perturbed with Gaussian noise and lie on the surface of a hypersphere. This was chosen in order to have highly non-linear data.

Mathematically, let **c**_*j*_ be the center of the *j*-th cluster sampled from **c**_*j*_ *∼ ℕ* (0, *I*). Each center is then normalized and scaled to lie on a hypersphere of radius *r*:

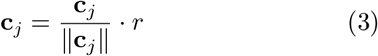

and thus ensuring that all cluster centers are equidistant from the origin. The data points for each cluster are generated by adding Gaussian noise *ϵ* to the cluster center:

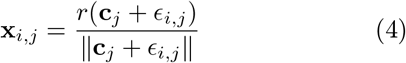

where:

- **x**_*i,j*_ is the *i*-th sample from the *j*-th cluster,
- *ϵ*_*i,j*_ *∼ ℕ* (0, *σ*^2^*I*) is the Gaussian noise with standard deviation *σ*

Again, normalization ensures that the points remain on the surface of a hypersphere of radius *r*. Finally, the data are standardized to have zero mean and unit variance, to have all the features on the same scale for model training.

Synthetic data are generated to form 10 clusters, varying the following parameters:

- Number of samples *N* : {50, 100, 150, 200, 500, 1000, 2000}.
- Number of features *d*: {100000, 200000, 500000}.
- Standard deviation of points *std*_*dev*_: {0.1, 1}
- Radius of the hypersphere: 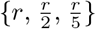, where 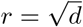

### 3.2 Models

Experiments were conducted on two generative models, the Deep Generative Decoder (DGD, [11]) and the Variational Autoencoder (VAE). For both models, a simple architecture was chosen for the decoder, involving only one hidden layer with dimension *h*_*dim*_ with a RELU activation function, a fixed latent space dimension *z*_*dim*_ = 16 and a single Gaussian prior with mean 0 and variance 1 on the latent space. The VAE encoder was a mirrored decoder. An Adam optimizer was chosen with a learning rate set to 10^*−*4^.

Both models were trained using early stopping on the validation set, with a patience of 10 epochs, and were subsequently evaluated on the test set.

In our experiments, the training size varies, while the validation set remains fixed at 400 samples and the test set at 1000 samples. These set sizes result from generating 5000 samples, applying an 80-20 train-test split, and then further splitting the training portion into training and validation sets using a 90-10 strategy. From the remaining training data, we sampled the desired number of instances for our experiments. Throughout all splits, the proportions of cluster labels are maintained. Five runs, for each set of parameters, were conducted for both models to assess performance variability.

Clusters were derived from the latent space using K-Means clustering.

## 4 Receptive field

In this section, we briefly introduce the concept of the receptive field, which was adopted in our experiments involving high-dimensional genomic data derived from both germline variants in the 1000 Genomes Project [12] and somatic variants from the International Cancer Genome Consortium (ICGC) database [13]. In these settings, where the number of features ranges from hundreds of thousands to several millions, a fully connected output layer would be computationally infeasible due to the extreme dimensionality of the data. The data under consideration are ordered for each chromosome, so in order to inform the network about the ordering, we are using a “receptive field” layer for the output. A fully connected output layer would also be challenging for the hardware with so many outputs.

The “receptive field” layer establishes connections between nodes in consecutive layers in a structured manner. Specifically, each node *i* in the current layer *L* is connected to a receptive region of width *w* in the next layer. Adjacent nodes *I* − 1 and *i* + 1 share an overlapping region of size *r*, ensuring local contextual learning while maintaining sparsity.

In particular, the number of neurons in the receptive field layer is dictated by the receptive width *w* and the overlapping region *r* between neighbors (the stride is *s* = *w* − *r*). Let us consider an output layer of dimension *d*. The number of neurons *h* in the receptive field are given by:

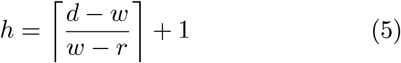

where ⌈*x*⌉ represents the “ceiling” function, which rounds *x* up to the nearest integer greater than or equal to *x*. This ensures that the number of neurons *h* is sufficient to cover the entire output dimension *d*.

The receptive field layer was applied both in the DGD and VAE models. For the DGD in the last layer of the decoder while for the VAE both in the last layer of the decoder and in the first layer of the encoder, using a transposed version.

A schematic example of the receptive field layer used in our application is shown in Figure 1.

**Figure 1:**
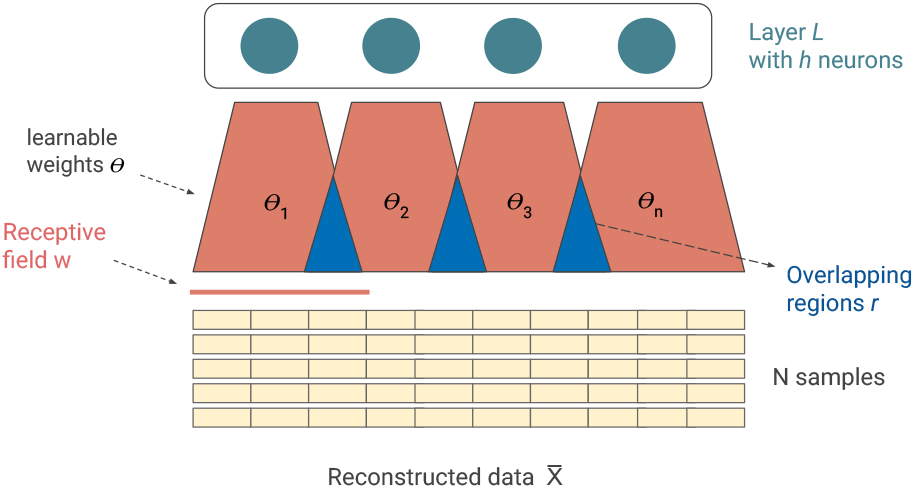
Receptive field layer with a receptive width *w* and overlapping region *r*. The layer connects layer with *h* neurons with the output data. Each neuron *i* has its own set of weights *θ*_*i*_.

## 5 Application to the 1000 Genome Project dataset

### 5.1 Dataset overview

In this study we used genomic variant data from the 1000 Genomes Project (1KGP) [12], one of the most comprehensive catalogs of human genetic variation across globally diverse populations. This international initiative sequenced over 2,500 individuals from 26 populations spanning Africa, Europe, East Asia, South Asia, and the Americas, producing high-coverage variant calls including SNPs, indels, and structural variants that capture inherited genetic diversity across continental groups. For this study, we used a curated 1KGP release obtained from [14] and stored in https://zenodo.org/records/14106454, consisting of whole-genome sequencing-derived genotype calls formatted for downstream analyses. The dataset comprises 2504 samples and includes variants filtered using a 5% minor allele frequency (MAF) threshold to retain common polymorphisms and reduce data sparsity. After preprocessing, the resulting dataset contained 6,864,700 SNPs, with values (0, 1, 2) representing allele dosage counts corresponding to the number of alternative alleles relative to the reference allele. Figure 2 shows the distribution of the five continental population in the dataset. To evaluate how training performance scales with data size, we constructed a series of subsampled datasets by varying both the number of samples and features. We started from the downsampled version of the 1KGP dataset from [14] which contains 686,471 SNPs and we use stratified sampling based on population labels, yielding dataset sizes of 100, 200, 500, and 1,000.

**Figure 2:**
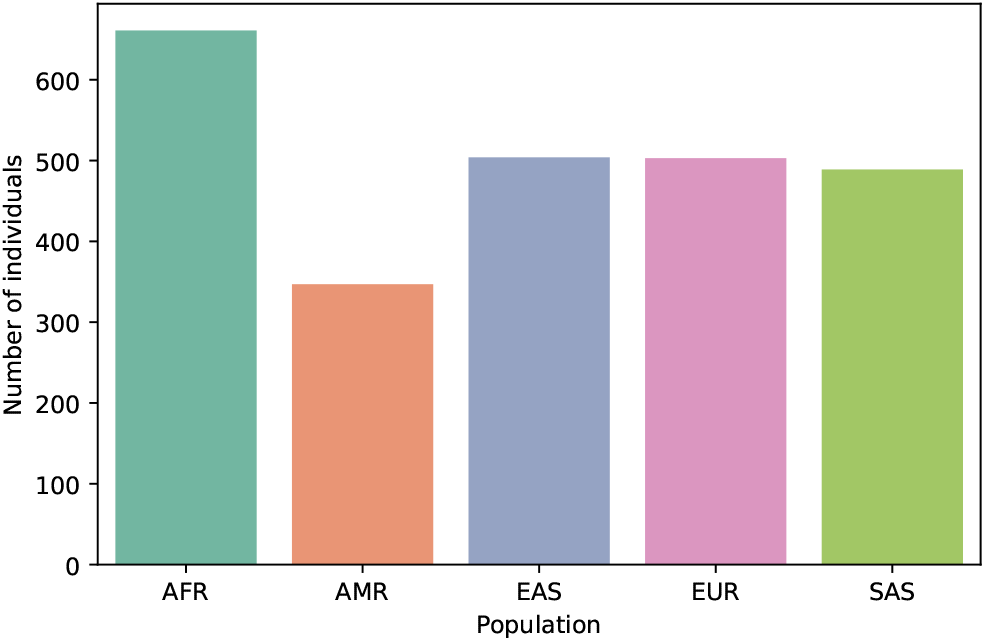
Distribution of individuals across the five continental population groups in the 1000 Genomes Project dataset used in this study. AFR: Africans; AMR: Admixed Americans; EAS: East Asians; EUR: Europeans; SAS: South Asians.

Feature subsets of 100,000, 200,000, and the 686,471 SNPs were considered. To account for variability introduced by random sampling, each configuration was repeated 3 times with different random seeds, producing independent subsampled datasets for each setting. In this way we assessed feature dimensionality and sample size impact on model performance, for both DGD and VAE models.

### 5.2 Models

For the DGD model, we set *z*_*dim*_ = 128, with information flowing through two hidden layers (256 and 512 neurons) with GELU activations and dropout rate of 0.2, followed by a final receptive field layer. The receptive field window size (*w*) was set to 800 with an overlap (*r*) of 200. The number of neurons in the receptive field layer depends on the feature set size, resulting in 167 (100k), 333 (200k), and 1,144 (680k) neurons for the three feature sizes, respectively. Since the decoder outputs a distribution over the three possible allelic states (0, 1, 2), a softmax activation is applied at the final layer. For the VAE, we used the same decoder architecture, while the encoder followed an inverse structure, compressing from the input space down to the latent space through symmetric receptive field blocks and two hidden layers. To handle the discrete nature of genotype input data (0, 1, 2), the encoder embeds each allelic value into a dense 32-dimensional vector before processing, allowing the model to learn a continuous representation of the discrete input. Additionally, the log-variance was clamped between −5 and 5 to ensure numerical stability during training. All models were trained with a learning rate of 1 × 10^*−*3^ with an Adam optimizer and a weight decay of 1 × 10^*−*5^ for a maximum of 300 epochs with early stopping on the test sets with a patience of 10 epochs. Batch sizes were adapted to the sample sizes used during training: 32 for datasets smaller than 200 samples, and 64 otherwise. A cross-entropy loss was used as the reconstruction error. No hyperparameter tuning was performed, and it is likely that better models could be found. Beyond the computational cost, systematic tuning via cross-validation was not practical given the limited number of samples in several configurations (as few as 100), which would leave too few held-out samples per population group. We emphasize that the goal of this paper is to analyze the feasibility of training in very high-dimensional settings with only few samples. For the same reason we do not use train-validation-test sets but rather only train and test sets, with a 80-20 split, to assess the presence of overfitting and to evaluate model performance. To evaluate model performance, we compare the cross-entropy loss across varying sample sizes and feature dimensionalities. Additionally, we assess the ability of both models to capture population structure by applying K-Means clustering on the learned latent representations, evaluating the resulting groups against ground-truth population labels using the Adjusted Rand Index (ARI) and Normalized Mutual Information (NMI).

## 6 Application to the ICGC dataset

### 6.1 Data preprocessing

The second dataset analyzed in this study was the 28th release of the International Cancer Genome Consortium (ICGC) database [13], downloaded on 03/05/2024. This release includes approximately 25,000 cancer samples across 86 cancer projects and 22 primary cancer sites.

Here, we focused exclusively on somatic single nucleotide variants (SNVs). To ensure data quality, we filtered the data set by selecting cancer samples with at least 100 SNVs and only including primary cancer types with a minimum of 80 samples.

In our experiments, we used a binary feature encoding: a value of 1 indicated the presence of a mutation at a given chromosome position, while 0 indicated its absence. We included chromosome positions only if at least two samples exhibited a mutation at that specific location. After applying this criterion, the final data set consisted of 5,176 samples and approximately 4.4 million genomic positions (features) across 22 cancer types.

While alternative filtering methods could have been used, we chose this approach to exclude cancer types with very few mutations and to maintain a robust sample size for each type. For chromosome position inclusion, we set the threshold at two samples to prevent excessive data reduction, given that our analysis operates at the chromosome positional level.

In Figure 3, we show the distribution of SNVs per tumor type in our dataset.

**Figure 3:**
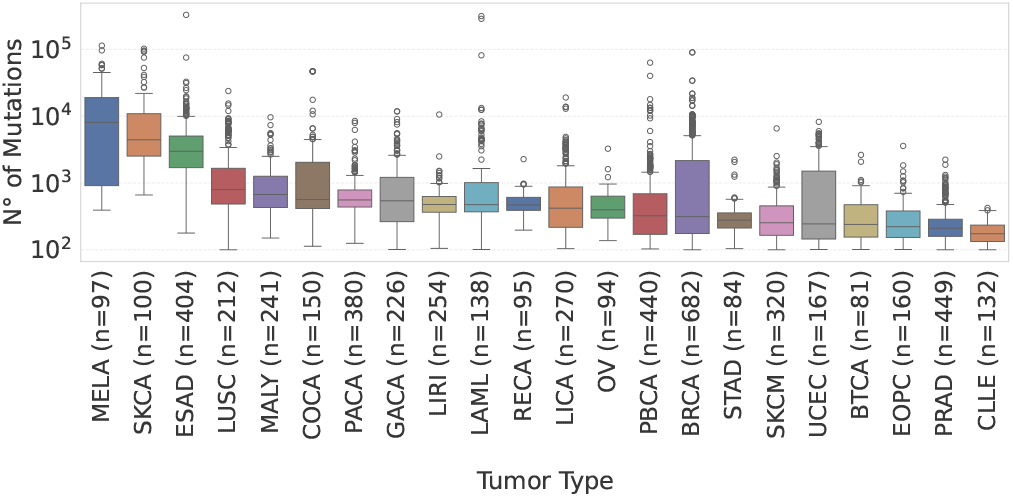
Distribution of the number of SNVs (log scale) for each tumor type on the filtered ICGC dataset. Abbreviations of cancer types can be found here https://gdc.cancer.gov/resources-tcga-users/tcga-code-tables/tcga-study-abbreviations.

### 6.2 Models

For the DGD model, we set *z*_*dim*_ to 128, followed by two hidden layers with GELU activation functions and a dropout rate of 0.2. Specifically, information flows from the latent space with 128 neurons through two layers of 256 and 512, followed by a final receptive field layer with 7,290 neurons which reconstruct the 4.4 million outputs with a sigmoid activation function.

The receptive field size (*w*) was set to 800, with an overlapping region (*r*) of 200.

For the VAE model, we used the same decoder architecture, while the encoder followed an inverse structure. In addition, for the VAE, we clamped the log-variance between -5 and 5 to prevent numerical instability during training. Both models were trained with a batch size of 80 and a learning rate of 1 × 10^*−*3^ with an Adam optimizer and a weight decay of 1 × 10^*−*5^ for a maximum of 500 epochs using early stopping with a patience of 10 epochs. During training, we applied a weighted binary cross-entropy loss for each output, assigning 100 times more weight to ones (mutation presence) than to zeros. This weighting was necessary to address the high sparsity of the data. No hyperparameter tuning was performed. Optimizing a model for such high-dimensional data would require a significant amount of time and computation, and, as in the 1KGP experiments, the limited number of samples per tumor type makes a further split for tuning purposes impractical. As highlighted in the previous experiment, we do not use train-validation-test sets but rather only train and test sets split to assess the presence of overfitting and to evaluate model performance. In particular we applied a 80-20 strategy, resulting in 4140 samples and 1036 for train and test sets, respectively Finally, we compared the DGD latent representation with VAE and PCA with the same number of components (*z*_*dim*_ = 128) for tumor type classification. Given the high dimensional dataset we use the Incremental PCA from the scikit-learn python package.

## 7 Results

### 7.1 Synthetic data

Figure 4 depicts the results for the Mean Squared Error (MSE) on synthetic data when varying the number of samples and features for both the DGD and VAE. The standard deviation of points is set to 0.1 and the radius to 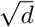, where *d* is the number of features. The plots show that the test MSE is consistently lower for the DGD than for the VAE for all feature dimensions and hidden dimensions (*h*_*dim*_).

**Figure 4:**
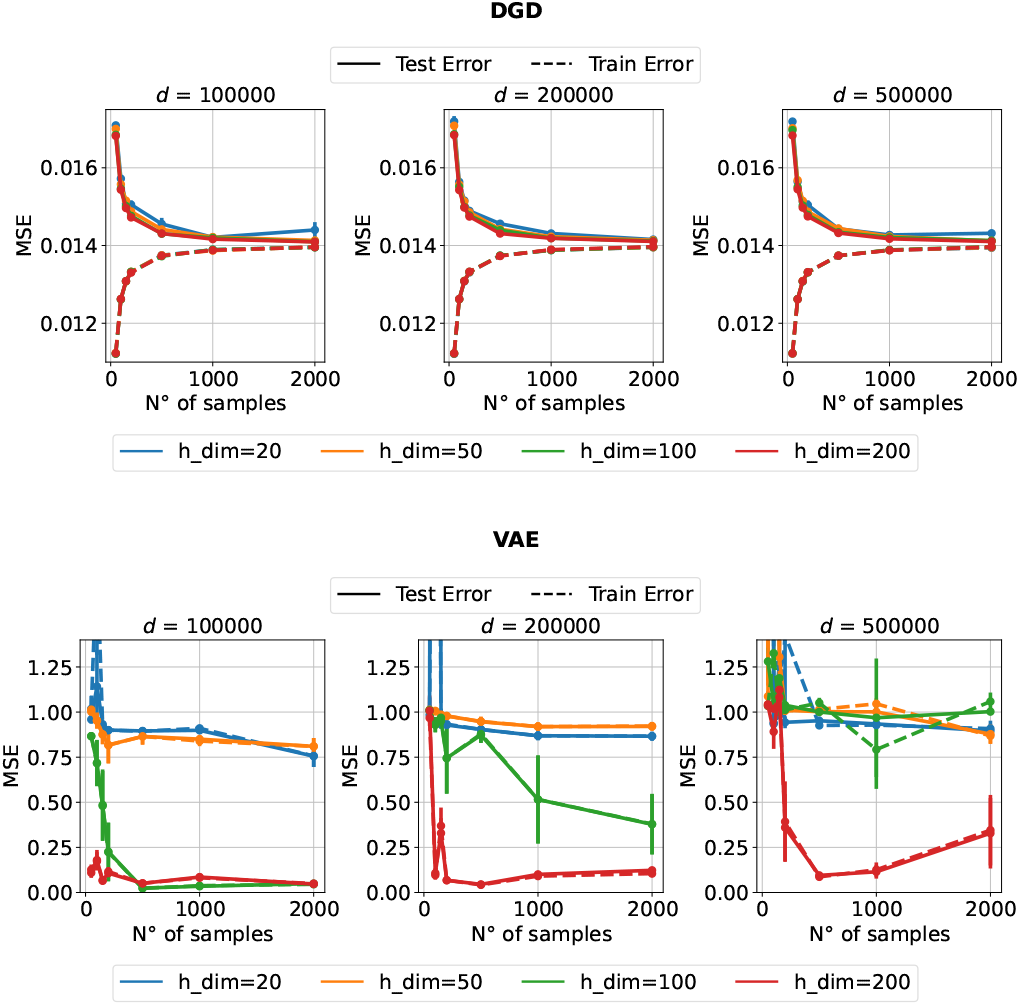
Synthetic data. Mean Squared Error (MSE) curves for the DGD (top) and VAE (bottom) for different numbers of samples and feature dimensions *d*. Dashed lines indicate the training set, while solid lines indicate the test set. Different colors correspond to different hidden layer dimensions *h*_*dim*_. Note the difference in scale between DGD and VAE.

Contrary to the DGD, the VAE performance is sensitive to the number of samples and hidden dimension *h*_*dim*_. It also exhibits unstable training behavior, although we clamped the log-variance to prevent numerical instability. To investigate whether this instability could be mitigated, we tested two additional training strategies: setting *β* = 0 and applying a linear warm-up of the KL divergence coefficient from 0 to 1 over the first 100 epochs, following previous approaches used to stabilize VAE training and mitigate posterior collapse [15]. The results, reported in Supplementary Figure S1, suggest that both strategies can improve the training behavior. In particular, reducing *β* or gradually increasing its contribution leads to lower MSE values for *h*_*dim*_ = 20, 50, and 100, and appears to partially stabilize the optimization process. However, the variance across different runs remains relatively large, especially for the smaller hidden dimensions (*h*_*dim*_ = 20 and 50), indicating that the instability is only partially mitigated. The primary aim of this work is not to compare the two models, which would have required a more complete optimization through hyperparameter tuning, but rather to study the feasibility of training in a high-dimensional settings. For the DGD, the MSE is almost independent of the feature dimension and depends solely on the number of samples as seen in figure 5. These results are consistent across all experiments, varying the standard deviation of points and radius of the hypersphere during data generation; see Supplementary Figures S2 and S3. This supports our hypothesis in Section 2.1 that the number of samples needed does not depend on the feature dimension. For the largest model (h-dim=200) this also holds for the VAE, but it seems much less stable. Regarding clustering performance both DGD and VAE can successfully recover clusters in the latent space with a K-Means algorithm, for all feature dimensionalities and hidden dimensions *h*_*dim*_.

**Figure 5:**
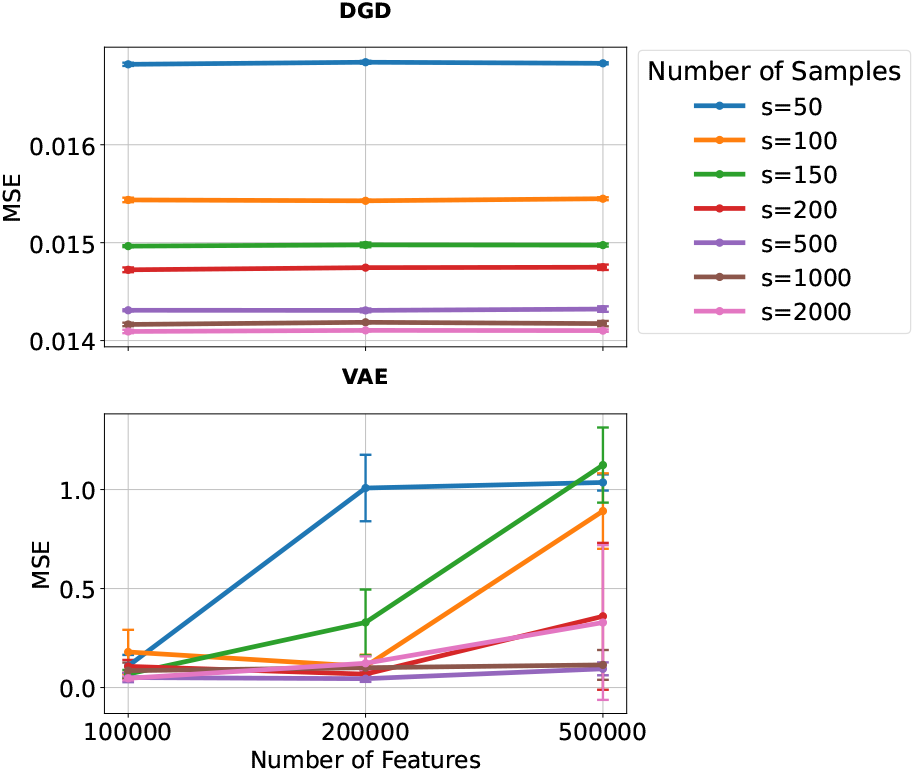
Synthetic data. Test error (MSE) when varying the number of features for different number of samples for hidden dimension *h*_*dim*_ = 200, for the DGD (top) and VAE (bottom). Note the different MSE scales in the two plots. Also note that the error bars are almost invisible in the DGD plot.

### 7.2 1KGP dataset

To test the proper functioning of the receptive field layer for both VAE and DGD models we perform preliminary experiments on the FashionMnist dataset. In particular, we train simple networks with *z*_*dim*_ = 32 for 50 epochs and evaluate them on the test set. In Supplementary Figures S4 and S5 we show the loss function and the PCA for the latent space for both the DGD and VAE models on the test sets. The results confirm that such an architecture is able to learn spatial information and to extract relevant structures in the data. We subsequently train both VAE and DGD for all sample sizes and feature dimensionalities for the 1KGP dataset. Figure 6 illustrates the cross-entropy loss for both DGD and VAE across all experiments. Both models exhibit the same trend: increasing the feature dimensionality has little effect on the Cross-Entropy Loss, which converges to similar values in each case. However, DGD consistently achieves lower cross-entropy loss than VAE across all sample sizes and feature dimensionalities.

**Figure 6:**
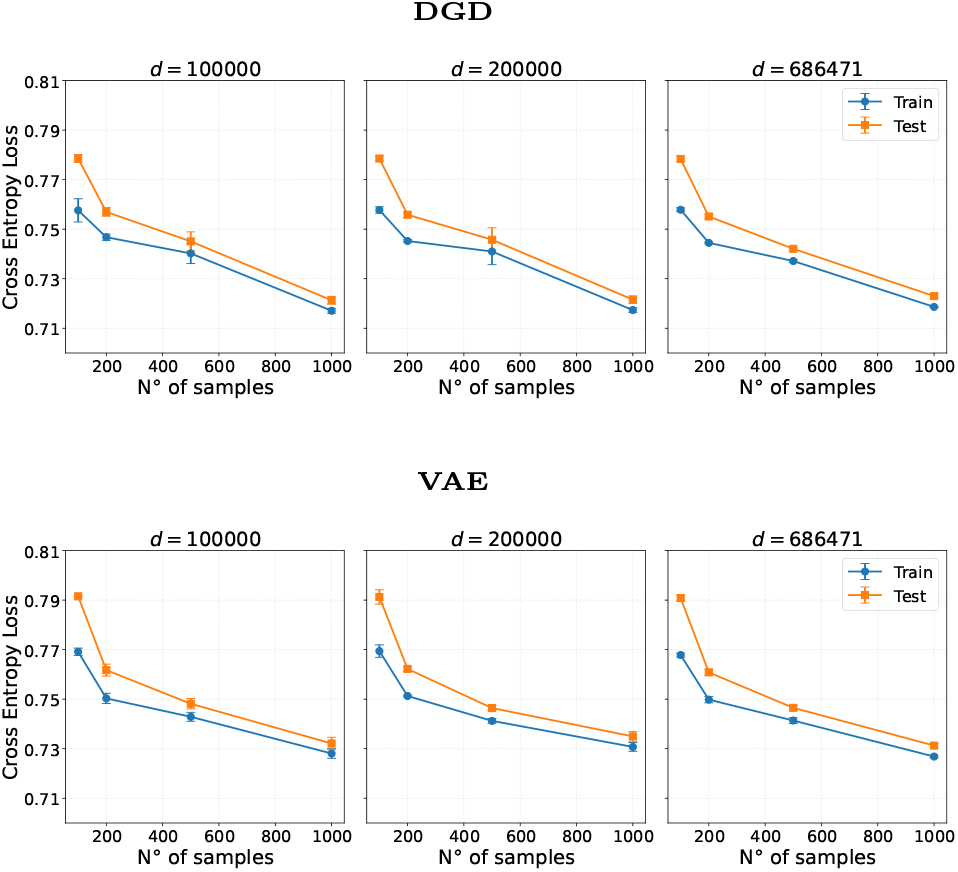
1KPG dataset. Mean cross entropy loss curves for the DGD (top figure) and VAE (bottom figure) for different number of samples and features *d*.

To further evaluate training feasibility in this high-dimensional setting, Table 1 reports the NMI and ARI scores obtained by applying K-means clustering to the 128-dimensional latent representations learned by DGD and VAE. NMI and ARI were computed using the available population information, and the number of clusters in K-means was therefore set to 5. For comparison, Table 1 also includes results obtained using PCA on the data normalize by allele frequency. In this case, we used 64 components for datasets with fewer than 100 samples, and 128 components for all larger datasets.

**Table 1:** NMI and ARI scores on the 1000 Genomes Project dataset for DGD, VAE, and PCA baselines across different training sizes and feature dimensionalities. Results are averaged over 3 runs (mean *±* std).

| Model | Size | $d = 100k$ | | $d = 200k$ | | $d = 680k$ | |
| --- | --- | --- | --- | --- | --- | --- | --- |
|  |  | NMI | ARI | NMI | ARI | NMI | ARI |
| DGD | 100 | 0.70±0.07 | 0.40±0.07 | 0.74±0.10 | 0.49±0.18 | 0.74±0.10 | 0.48±0.19 |
|  | 200 | 0.71±0.07 | 0.59±0.08 | 0.76±0.08 | 0.60±0.09 | 0.73±0.07 | 0.57±0.07 |
|  | 500 | 0.73±0.06 | 0.61±0.03 | 0.74±0.05 | 0.62±0.01 | 0.75±0.03 | 0.62±0.06 |
|  | 1000 | 0.76±0.04 | 0.69±0.02 | 0.75±0.03 | 0.63±0.05 | 0.87±0.04 | 0.85±0.05 |
| VAE | 100 | 0.51±0.09 | 0.18±0.10 | 0.62±0.06 | 0.37±0.05 | 0.62±0.05 | 0.34±0.06 |
|  | 200 | 0.53±0.04 | 0.37±0.07 | 0.54±0.05 | 0.39±0.05 | 0.72±0.05 | 0.58±0.04 |
|  | 500 | 0.65±0.05 | 0.49±0.05 | 0.64±0.02 | 0.53±0.05 | 0.81±0.03 | 0.79±0.02 |
|  | 1000 | 0.68±0.06 | 0.55±0.09 | 0.63±0.02 | 0.52±0.04 | 0.84±0.05 | 0.85±0.08 |
| PCA | 100 | 0.95±0.05 | 0.91±0.10 | 0.95±0.05 | 0.91±0.10 | 0.95±0.05 | 0.91±0.10 |
|  | 200 | 0.91±0.02 | 0.86±0.02 | 0.91±0.02 | 0.86±0.02 | 0.91±0.02 | 0.86±0.02 |
|  | 500 | 0.93±0.01 | 0.90±0.02 | 0.93±0.01 | 0.90±0.02 | 0.93±0.01 | 0.90±0.02 |
|  | 1000 | 0.92±0.02 | 0.89±0.03 | 0.92±0.02 | 0.89±0.03 | 0.92±0.02 | 0.89±0.03 |

Overall, DGD consistently outperforms VAE across most configurations, particularly at lower dimensionalities, suggesting a better ability to capture the underlying cluster structure. Both models show a clear trend of improving performance as the feature dimensionality increases, with the most notable gains observed at 1000 samples on the 680*k* dataset, where DGD reaches *NMI* = 0.87 and *ARI* = 0.85, and VAE achieves *NMI* = 0.84 and *ARI* = 0.85. Interestingly, the performance gap between the two models narrows at higher dimensionalities and larger sample sizes, indicating that VAE benefits more from increased data and feature richness. At lower dimensionalities (Size=100, 200), DGD holds a more substantial advantage, especially in *ARI*, which is more sensitive to cluster assignment quality. However, PCA appears to yield stronger clustering performance in this setting, which may indicate that the main sources of variation in the genotype data are more readily captured by a linear representation than by the VAE and DGD latent spaces. In addition, since we did not perform hyperparameter tuning, the chosen model settings may not be optimal and could have affected the observed performance. Furthermore, it is important to note that both VAE and DGD are primarily optimized for reconstruction tasks; therefore, improvements in reconstruction quality do not necessarily translate into optimal clustering structure in the learned latent representations. Notably, with 1,000 samples and the maximum number of features (680*k*), clustering performance is comparable across methods. Figure 7 illustrates the corresponding latent representations for all three approaches. Nevertheless, the results support our hypothesis that it is possible to successfully train generative models in high-dimensional settings even with a limited number of training samples.

**Figure 7:**
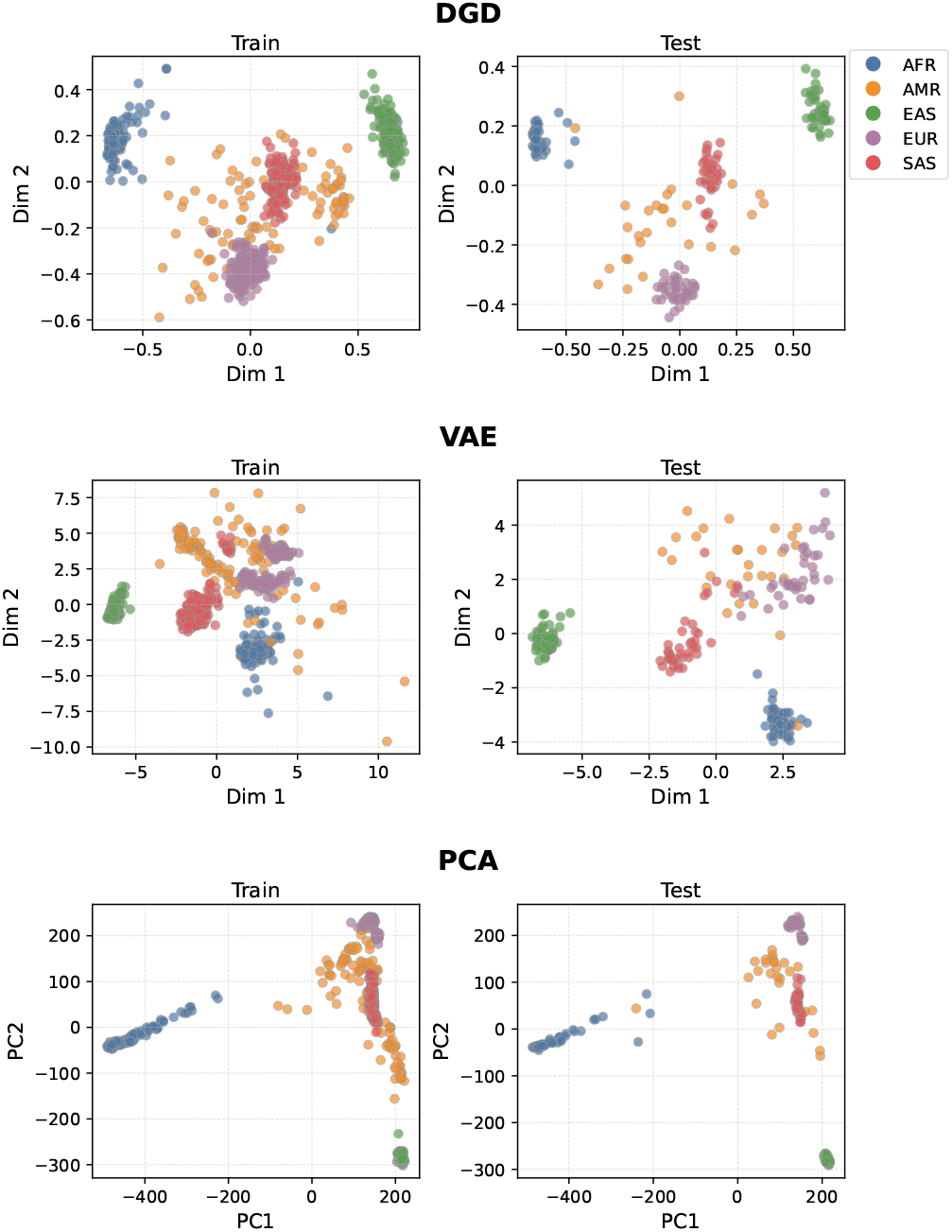
1KGP dataset. A 2d PCA representation computed for both the training and test sets for the DGD, VAE, and PCA models, using 1,000 samples and 680k features

To further investigate the contribution of individual population groups to the clustering structure, we performed an additional ablation experiment by removing the Admixed Americans (AMR) population. By definition, AMR individuals carry ancestry from multiple continental populations, including European, African, and Native American groups, which causes them to occupy intermediate positions between continental clusters in low-dimensional representations rather than forming a distinct group. This admixture pattern is therefore expected to introduce ambiguity in the learned latent representations, potentially influencing cluster separability. Results are reported in Supplementary Table S1.

When AMR samples are excluded, DGD shows consistent and robust performance across all configurations, with NMI stabilizing around 0.82–0.84 already at low sample sizes and across all feature dimensionalities, confirming its ability to capture population structure even with limited data. In particular, ARI reaches 0.97 at 1,000 samples on the 680k dataset. Importantly, DGD achieves strong clustering already at *d* = 100k suggesting that it does not require high feature dimensionality to learn meaningful representations. VAE, in contrast, requires both larger sample sizes and higher dimensionality to reach comparable performance.

### 7.3 ICGC dataset

We train DGD and VAE models on the ICGC dataset using the receptive field layer. Figure 8 shows the complete training and test losses for both models. The minimum BCE values on the test sets were 1.57 × 10^4^ for DGD and 1.67 × 10^4^ for VAE, respectively.

**Figure 8:**
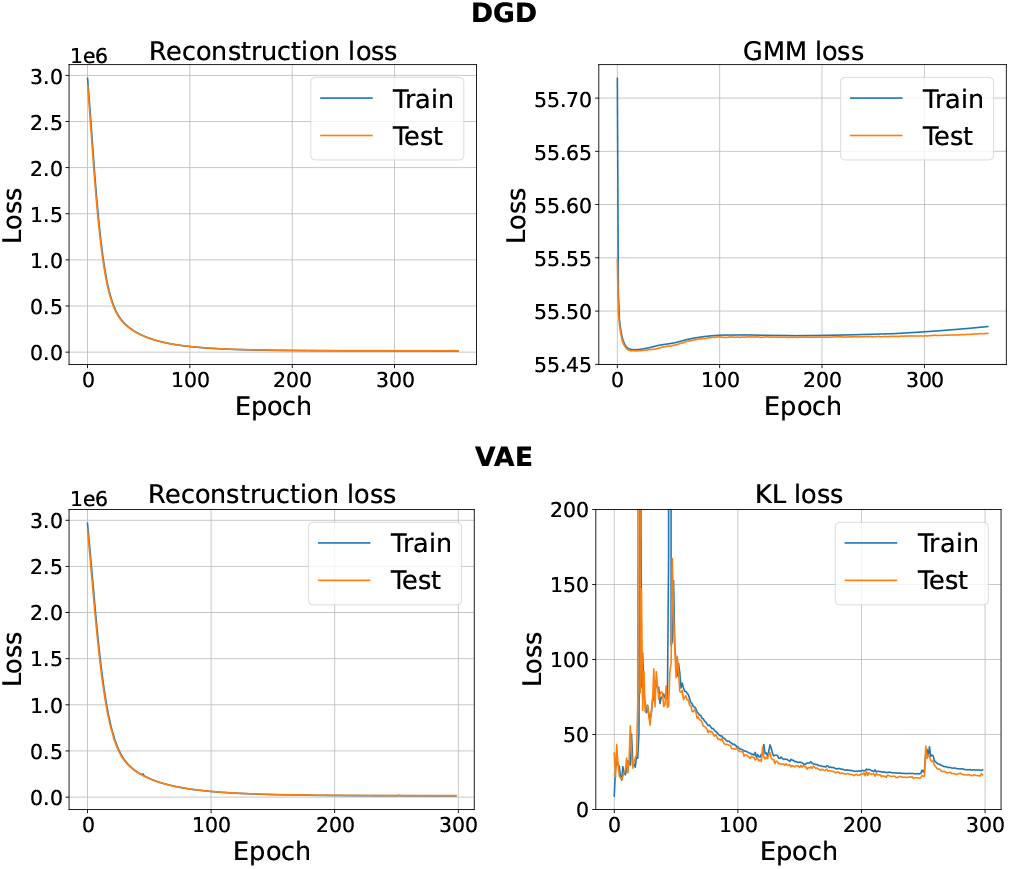
ICGC data. Reconstruction loss (BCE) together with the GMM loss for the DGD (top) and KL divergence loss for the VAE model (bottom). For both figures, training losses are represented by blue lines and test losses by orange ones.

To further evaluate model performance, we begin by examining the latent spaces produced by both models. Interestingly, the latent space of the VAE appears less organized compared to that of the DGD, which has evidently captured tumor-specific structures, as illustrated in Figure 9.

**Figure 9:**
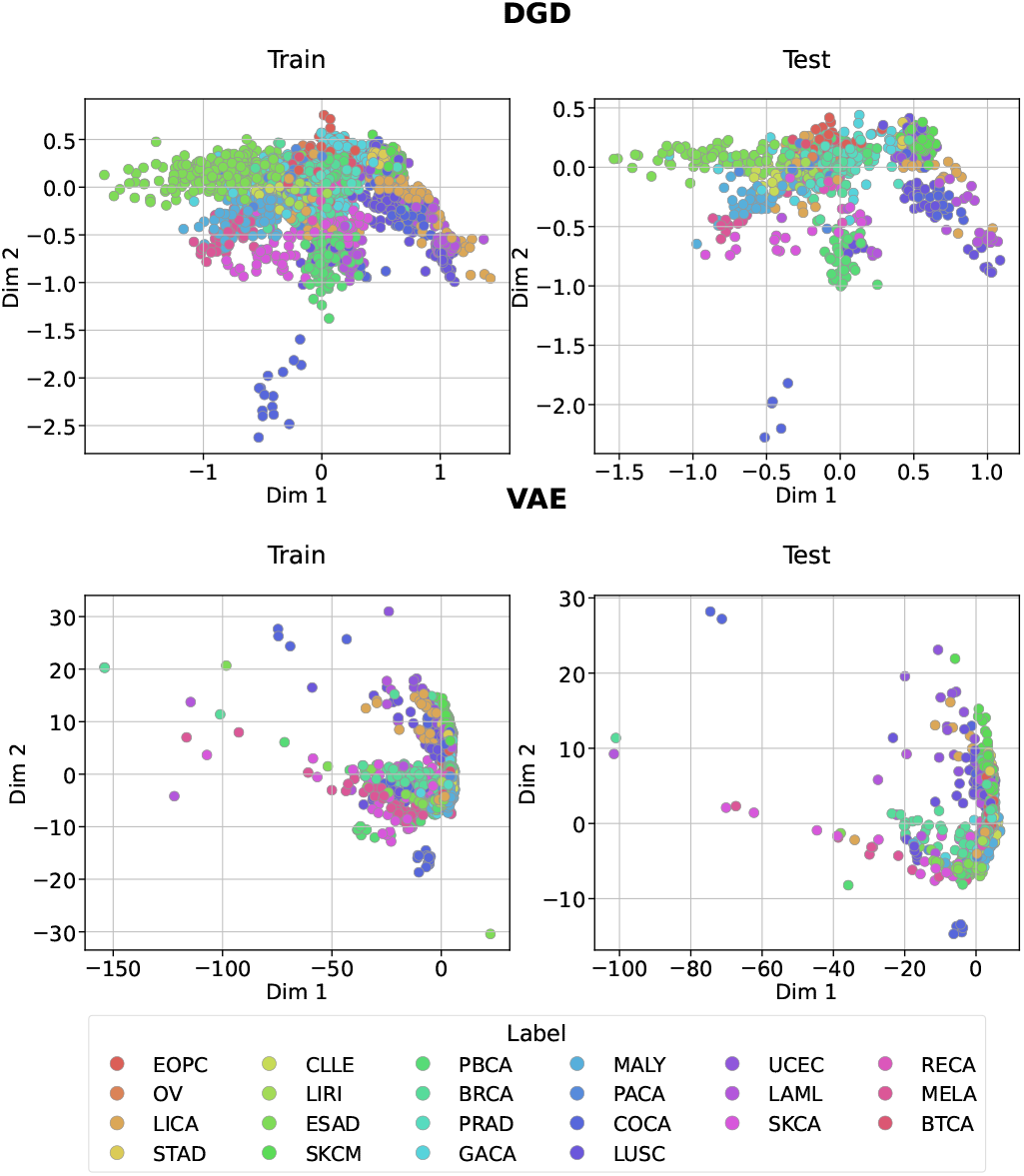
PCA of the latent spaces generated by the DGD (top) and VAE (bottom) for ICGC train and test data. Colored by cancer type.

In Figure 10, we also show the PCA directly performed on the raw ICGC dataset for a qualitative comparison with the DGD and VAE representations. To quantitatively assess performance, we first evaluate the models by applying K-Means clustering to the latent representations (*z*_*dim*_ = 128), using 22 clusters corresponding to the number of cancer types. We then compute ARI and NMI for DGD, VAE, and PCA. DGD achieves the best performance (ARI: 0.53, NMI: 0.25), followed by VAE (0.44, 0.16) and PCA (0.16, 0.01)

**Figure 10:**
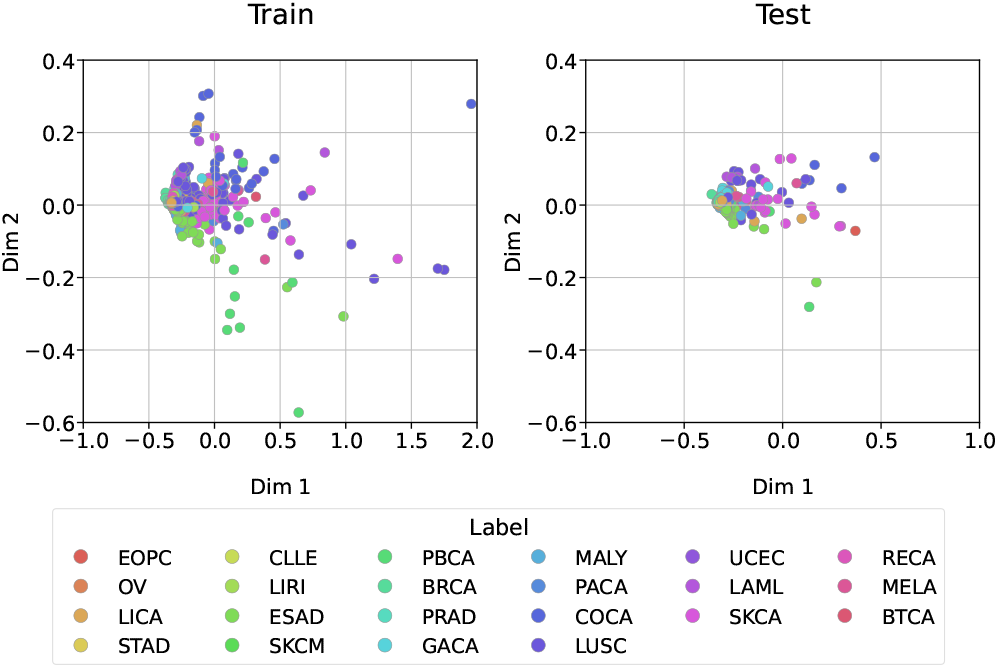
PCA performed on the ICGC dataset.

Furthermore we evaluated models on a tumor classification task using the latent representation with *z*_*dim*_ = 128. The task involved classifying 22 distinct tumor types. We trained a balanced logistic regression model using latent representations from the training set and evaluated performance on the latent representations of the test set. To assess the robustness of the classification results, 95% confidence intervals for balanced accuracy were computed via bootstrap resampling of the test set predictions (1,000 iterations).

To determine whether DGD offers advantages in low-dimensional representation, we compared the classification results against VAE representations and also with those obtained from PCA with equivalent 128 dimensions. Figure 11 presents confusion matrices for the three approaches. Our analysis reveals that DGD outperforms VAE in all tumour types and PCA in 15 of 22 tumour types. The overall balanced accuracies of DGD, VAE, and PCA are 67% (95% CI: 64–70%), 42% (95% CI: 39–45%), and 54% (95% CI: 51–57%), respectively.

**Figure 11:**
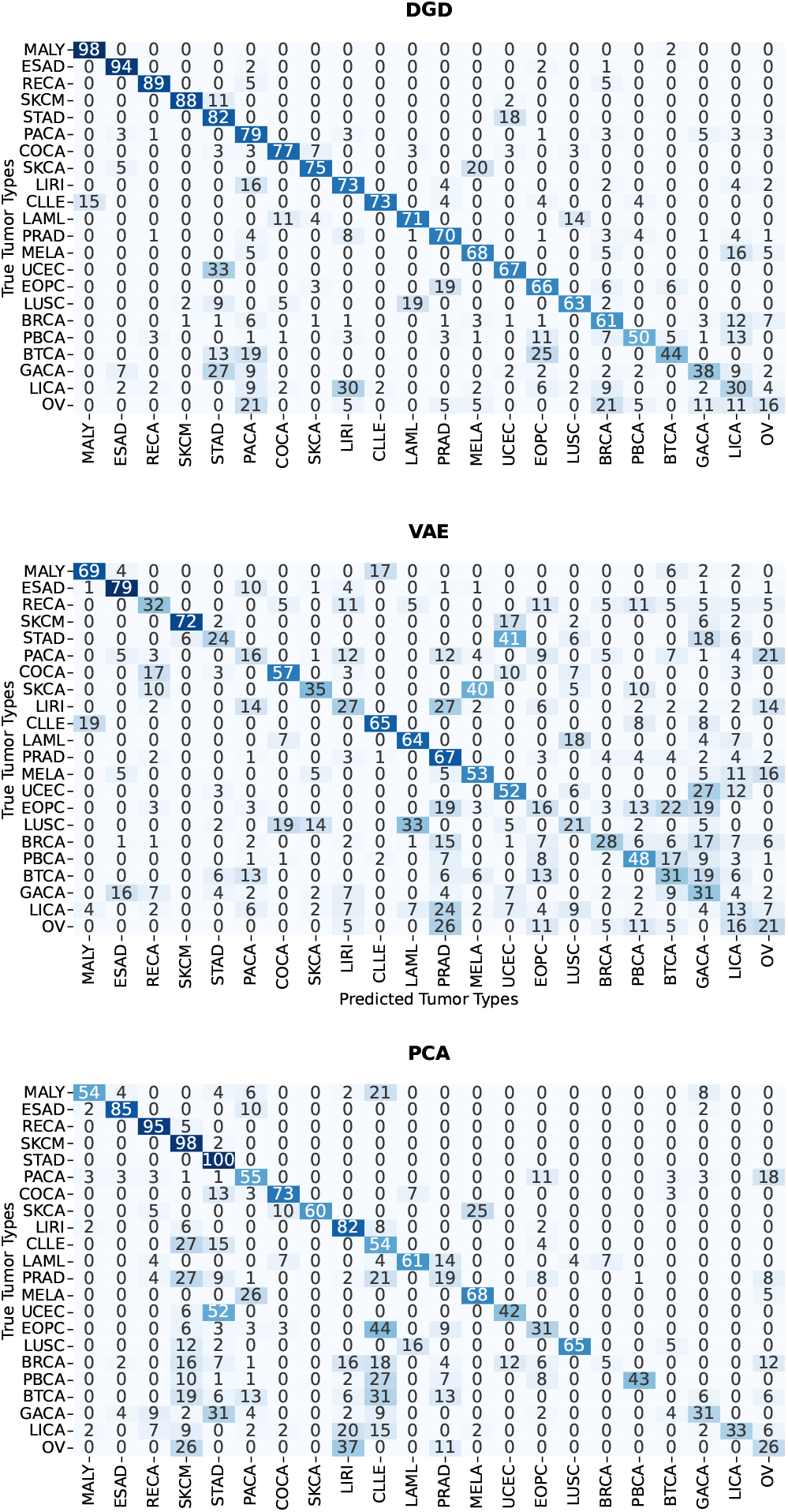
Confusion matrices illustrating the classification of tumor types based on the latent space representations. Numbers represents percentages. The results from DGD are shown at the top, VAE in the middle while the PCA results are displayed at the bottom

Among the seven tumor types where PCA demonstrates better performance compared to the DGD, five show very similar results.

Notably, for Stomach Adenocarcinoma (STAD) and Liver Cancer (LIRI), PCA effectively captures data relationships, despite high false positives rate, while the DGD exhibits lower performance.

However, the DGD significantly outperforms PCA for several tumor types where PCA largely fails, in particular for Malignant Lymphoma (MALY), Pancreatic Cancer (PACA), Biliary Tract Carcinoma (BTCA), Breast Cancer (BRCA), Prostate Adeno Carcinoma (PRAD), Early Onset Pancreatic Cancer (EOPC) and others. These performance differences can be attributed to the DGD ability to model nonlinear processes, which PCA cannot effectively capture.

To assess whether the VAE’s underperformance is sensitive to the choice of the KL weighting term *β*, rather than reflecting an intrinsic architectural limitation, we repeated training with *β* = 0 and with a linear KL warm-up (as in the synthetic data experiments). Table 2 reports balanced accuracy, macro precision and macro F1 for these variants alongside the models already discussed. Setting *β* = 0 substantially improves VAE performance (balanced accuracy 0.52, 95% CI 0.49–0.55), bringing it level with PCA and in fact surpassing it on macro F1 (0.52 vs. 0.49), while KL warm-up yields a smaller improvement (0.44, 95% CI 0.41–0.48) over the default VAE (*β* = 1, 0.42). PCA shows the widest gap between macro precision (0.58) and macro F1 (0.49), a pattern consistent with confident but infrequent predictions on a subset of classes. Despite these improvements, DGD retains a substantial advantage (0.67, 95% CI 0.64–0.70) and is the only model whose macro precision and recall are closely matched (0.66 vs. 0.67, the latter equal to balanced accuracy), suggesting balanced performance across tumor types. Full latent-space visualizations and confusion matrices for the *β* = 0 and linear warm-up variants are reported in Supplementary Figure S6 and Figure S7.

**Table 2:** Classification performance (95% CI) for ICGC tumor type classification, including the VAE *β* ablation. Precision and F1 are macro-averaged over tumor types; macro recall is by definition identical to balanced accuracy and is therefore not reported separately.

| Model | Balanced Acc. | Precision | F1 |
| --- | --- | --- | --- |
| DGD | 0.67 (0.64–0.70) | 0.66 (0.63–0.69) | 0.65 (0.62–0.68) |
| VAE ($\beta = 1$) | 0.42 (0.39–0.45) | 0.42 (0.40–0.46) | 0.40 (0.37–0.43) |
| VAE ($\beta = 0$) | 0.52 (0.49–0.55) | 0.56 (0.53–0.59) | 0.52 (0.48–0.54) |
| VAE ($\beta$ warm-up) | 0.44 (0.41–0.48) | 0.47 (0.43–0.50) | 0.42 (0.39–0.45) |
| PCA | 0.54 (0.51–0.57) | 0.58 (0.55–0.60) | 0.49 (0.46–0.52) |

Our findings demonstrate that training a decoder-only model with millions of features on a relatively small sample set (a few thousand samples) is not only feasible, but potentially advantageous for downstream applications, because it produces better separation among clusters in the latent space, an advantage that persists even after adjusting the VAE’s *β* parameter.

Furthermore, the DGD offers substantial computational efficiency advantages. Processing millions of features with PCA imposes significant computational demands. Even with incremental PCA (via scikit-learn), processing millions of features is memory-intensive. In contrast, we successfully trained DGD using a T4 GPU with 16 GB of memory, highlighting its remarkable efficiency for high-dimensional data processing.

## 8 Discussion

In this study, we hypothesize that training a decoder-only model is feasible in high-dimensional spaces and that the number of samples required to train such a model is independent of feature dimensionality and depends solely on the number of neurons in the network. Through experiments with synthetic data, we demonstrate that, across various settings and scenarios, the Mean Squared Error (MSE) for the Deep Generative Decoder (DGD) model remains effectively independent of feature dimensionality, whereas a Variational Autoencoder (VAE) does not exhibit this property.

To validate these findings on real genomic data, we further evaluated both DGD and VAE on the 1000 Genomes Project (1KGP) dataset, a large-scale human genetic variation resource comprising samples from diverse populations worldwide. We trained both models across varying feature dimensionalities and dataset sizes, assessing the quality of the learned latent representations through clustering metrics and reconstruction quality. Our results confirm that DGD consistently produces more structured latent representations than VAE, better capturing population structure across different configurations. Furthermore, both models exhibit comparable reconstruction quality, with DGD achieving marginally better performance, and neither model shows signs of overfitting even in high-dimensional regimes.

To further investigate this, we implemented a DGD and trained it on 4.4 million features derived from the Single Nucleotide Variants (SNVs) of the ICGC dataset across 22 cancer types. To allow the network to learn neighborhood relations and still pick up long-range correlations, we incorporated a receptive field layer into the decoder. Our results show that this model can be trained with a few thousands of samples and produces a useful, structured latent representation, where cancer types are well-clustered. In contrast, the VAE (with default *β* = 1) exhibits poor structure in its latent space; an ablation with *β* = 0 partially reduces this gap, suggesting that the choice of *β* contributes to, but does not fully explain, the VAE’s lower performance. We compared the latent representation generated by the DGD model with that obtained from Principal Component Analysis (PCA) using the same number of components. Through a classification task, we demonstrated that the DGD model outperforms both VAE and PCA in terms of tumor type classification accuracy. Additionally, our model offers an efficient alternative since it can be trained on a T4 GPU, requiring less RAM than PCA, especially in high-dimensional spaces.

Despite these results, a limitation of this work is that, given the computational cost of training on millions of features and the limited sample sizes available, we did not perform systematic hyperparameter tuning for either DGD or VAE. The reported comparisons should therefore be interpreted as evidence of feasibility in this high-dimensional, sample-limited regime, rather than as an optimized benchmark of the two architectures. Our targeted ablation on the VAE’s *β* parameter indicates that at least part of the observed gap is attributable to this specific hyperparameter choice rather than to an intrinsic limitation of the encoder-decoder architecture, and a more exhaustive tuning of both models could narrow the gap further. For the same reason, early stopping was based on the test set rather than a separate validation set, since further splitting the already limited training data risked producing an unstable stopping criterion; as a consequence, the reported test losses and downstream clustering and classification metrics should be interpreted as optimistic estimates of generalization performance rather than fully unbiased ones.

