## Supplementary materials for "A generative model for dimensionality reduction with millions of features and few samples"

#### 1 Synthetic data

In Figure S1, we present additional experiments under the setting $std_{dev} = 0.1$ and $r = \sqrt{d}$ for VAE. Panel (a) corresponds to the VAE model with $\beta = 0$, whereas panel (b) shows the results obtained by linearly increasing the KL divergence coefficient from 0 to 1 over the first 100 training epochs.

Figure S2 and S3 additional experiments for the DGD on synthetic data varying the number of samples and features with different combinations of standard deviation of points and hypersphere radius, are shown.

#### 2 Experiments on 1kgp

In Supplementary Table S1 we reporte NMI and ARI scores on the 1000 Genomes Project dataset after removing AMR samples, for DGD and VAE across different training sizes and feature dimensionalities.

#### 3 Experiments on FashionMnist

In Figures S4 we show the train and test losses for DGD and VAE with both receptive field architecture on FashionMnist dataset.

In Figure S5 we plot the PCA of the latent space colored by labels, here indicated for simplicity with numbers from zero to nine.

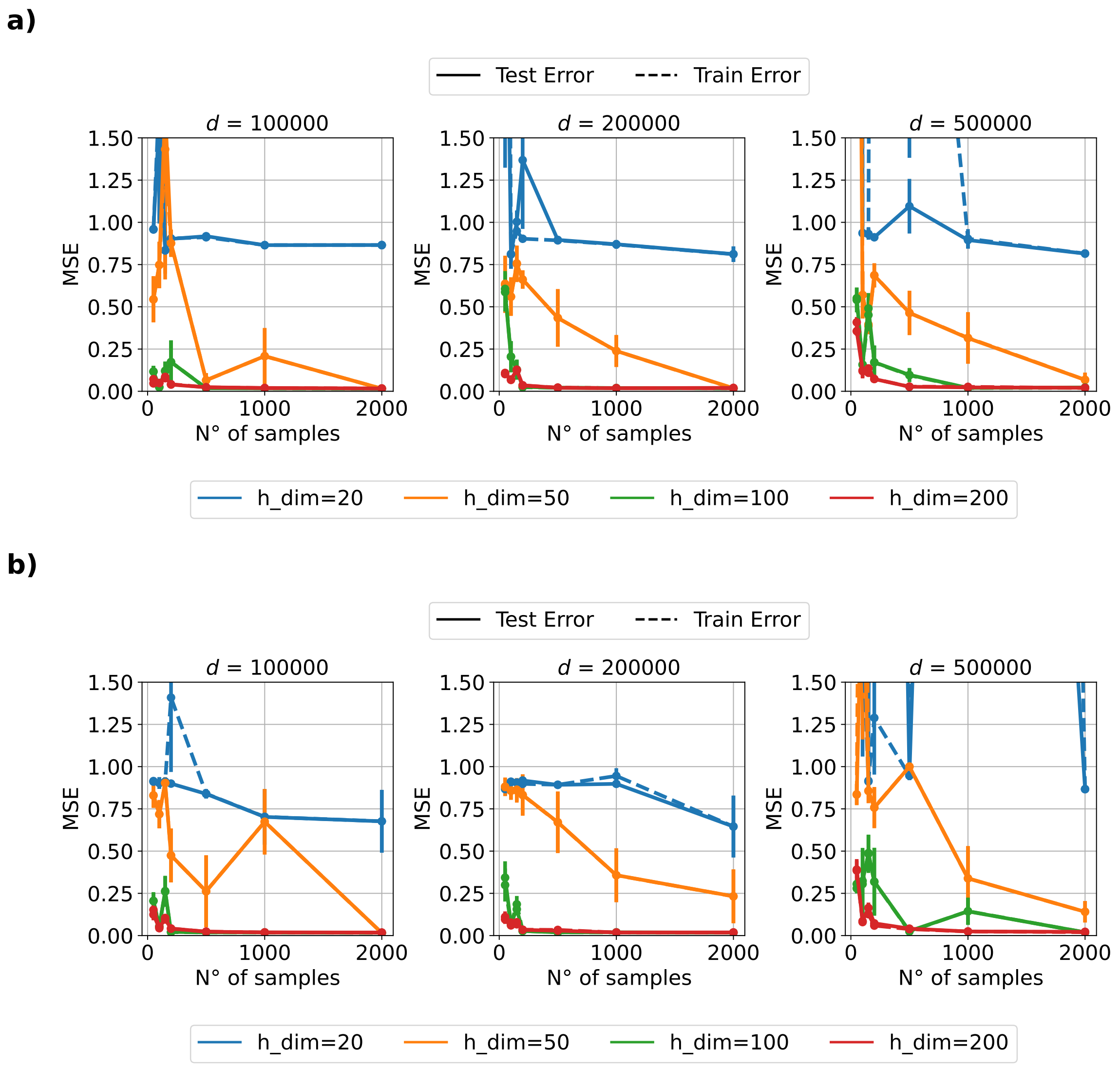


Figure S1: VAE experiments comparing different $\beta$ values. The top panel corresponds to the linear warm-up strategy, while the bottom panel uses fixed $\beta = 0$.

### 4 Experiments on ICGC

Figure S6 shows the two-dimensional principal component analysis (PCA) of the VAE latent representations. Panel (a) corresponds to the latent space learned by a VAE trained with $\beta = 0$, whereas panel (b) shows the latent space obtained using a KL-divergence warm-up strategy.

Figure S7 presents the confusion matrices obtained for the two VAE training strate-

| Model | Size | $d = 100\text{k}$ | | $d = 200\text{k}$ | | $d = 680\text{k}$ | |
|---|---|---|---|---|---|---|---|
| | | NMI | ARI | NMI | ARI | NMI | ARI |
| DGD | 100 | 0.82±0.01 | 0.60±0.03 | 0.82±0.09 | 0.67±0.17 | 0.82±0.10 | 0.65±0.19 |
| | 200 | 0.82±0.02 | 0.69±0.05 | 0.83±0.02 | 0.71±0.09 | 0.82±0.03 | 0.69±0.05 |
| | 500 | 0.82±0.03 | 0.71±0.03 | 0.83±0.02 | 0.69±0.04 | 0.83±0.03 | 0.73±0.02 |
| | 1000 | 0.82±0.02 | 0.75±0.11 | 0.84±0.04 | 0.71±0.08 | 0.98±0.06 | 0.97±0.06 |
| VAE | 100 | 0.57±0.05 | 0.35±0.05 | 0.62±0.09 | 0.39±0.13 | 0.70±0.12 | 0.49±0.10 |
| | 200 | 0.63±0.04 | 0.50±0.05 | 0.62±0.09 | 0.52±0.11 | 0.83±0.02 | 0.74±0.07 |
| | 500 | 0.80±0.03 | 0.69±0.06 | 0.74±0.02 | 0.69±0.02 | 0.98±0.03 | 0.98±0.03 |
| | 1000 | 0.84±0.06 | 0.74±0.15 | 0.74±0.05 | 0.69±0.08 | 0.94±0.04 | 0.95±0.04 |

Table S1: NMI and ARI scores on the 1000 Genomes Project dataset after removing AMR samples, for DGD and VAE across different training sizes and feature dimensionalities. Results are averaged over 3 runs (mean ± std).

gies. The top panel corresponds to the VAE trained with $\beta = 0$, whereas the bottom panel shows the results obtained using the KL-divergence warm-up strategy.

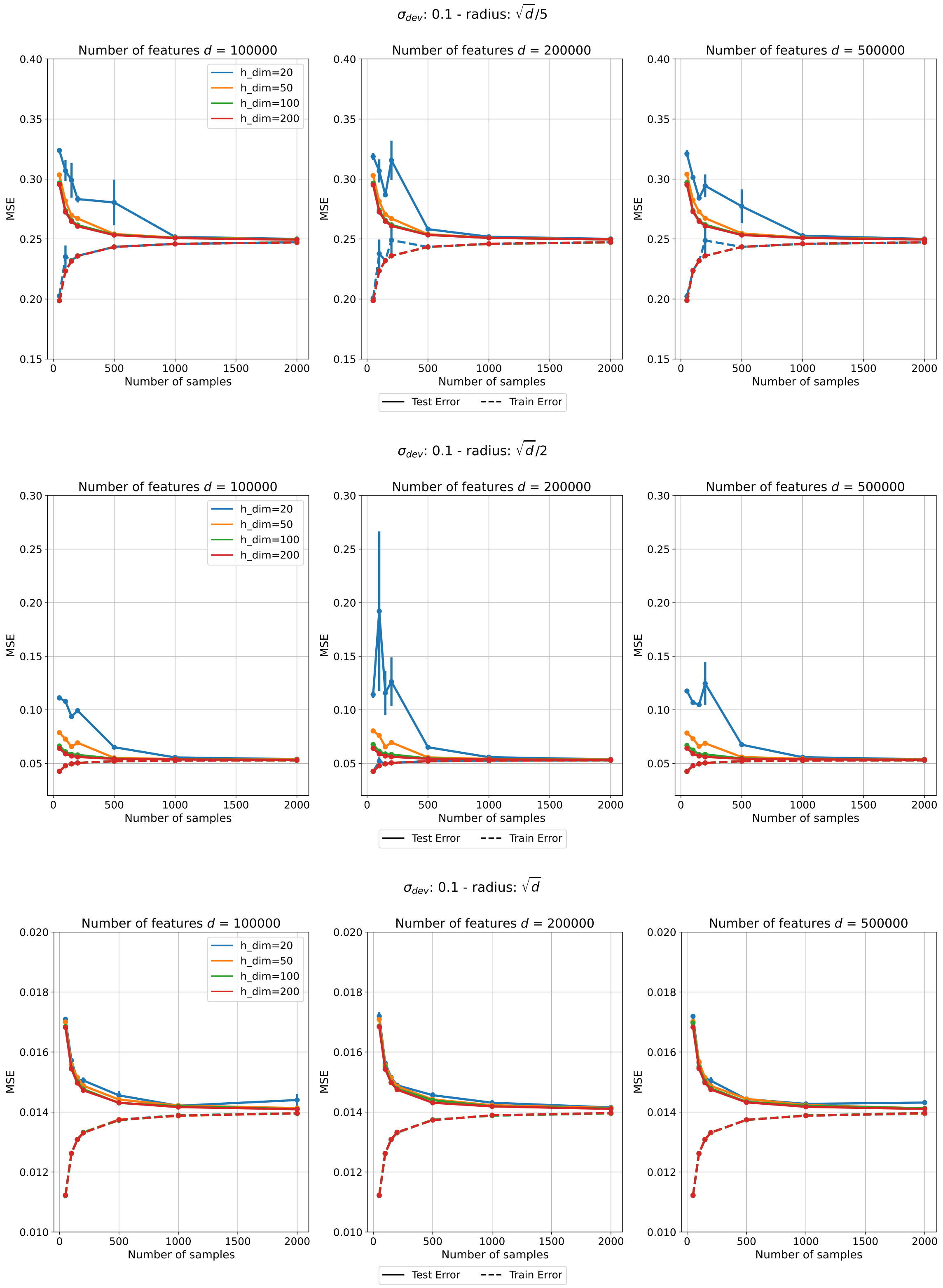
$\sigma_{dev}$: 0.1 - radius: $\sqrt{d}/5$
Number of features $d$ = 100000
Number of features $d$ = 200000
Number of features $d$ = 500000
h_dim=20
h_dim=50
h_dim=100
h_dim=200
MSE
Number of samples
Test Error
Train Error
$\sigma_{dev}$: 0.1 - radius: $\sqrt{d}/2$
Number of features $d$ = 100000
Number of features $d$ = 200000
Number of features $d$ = 500000
h_dim=20
h_dim=50
h_dim=100
h_dim=200
MSE
Number of samples
Test Error
Train Error
$\sigma_{dev}$: 0.1 - radius: $\sqrt{d}$
Number of features $d$ = 100000
Number of features $d$ = 200000
Number of features $d$ = 500000
h_dim=20
h_dim=50
h_dim=100
h_dim=200
MSE
Number of samples
Test Error
Train Error

Figure S2

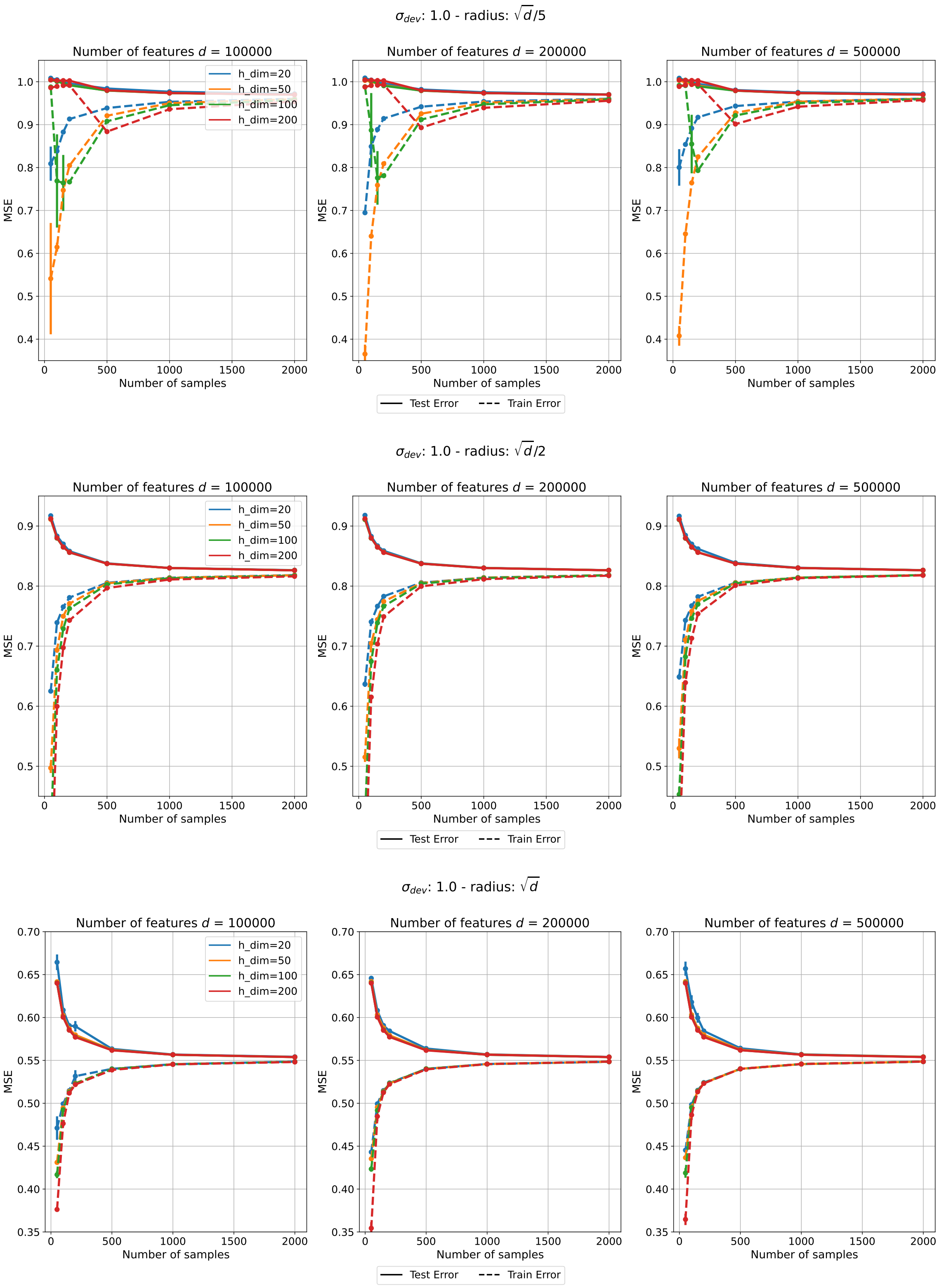

$\sigma_{dev}$: 1.0 - radius: $\sqrt{d}/5$
Number of features $d$ = 100000
Number of features $d$ = 200000
Number of features $d$ = 500000
h_dim=20
h_dim=50
h_dim=100
h_dim=200
MSE
Number of samples
Test Error
Train Error
$\sigma_{dev}$: 1.0 - radius: $\sqrt{d}/2$
$\sigma_{dev}$: 1.0 - radius: $\sqrt{d}$


Figure S3

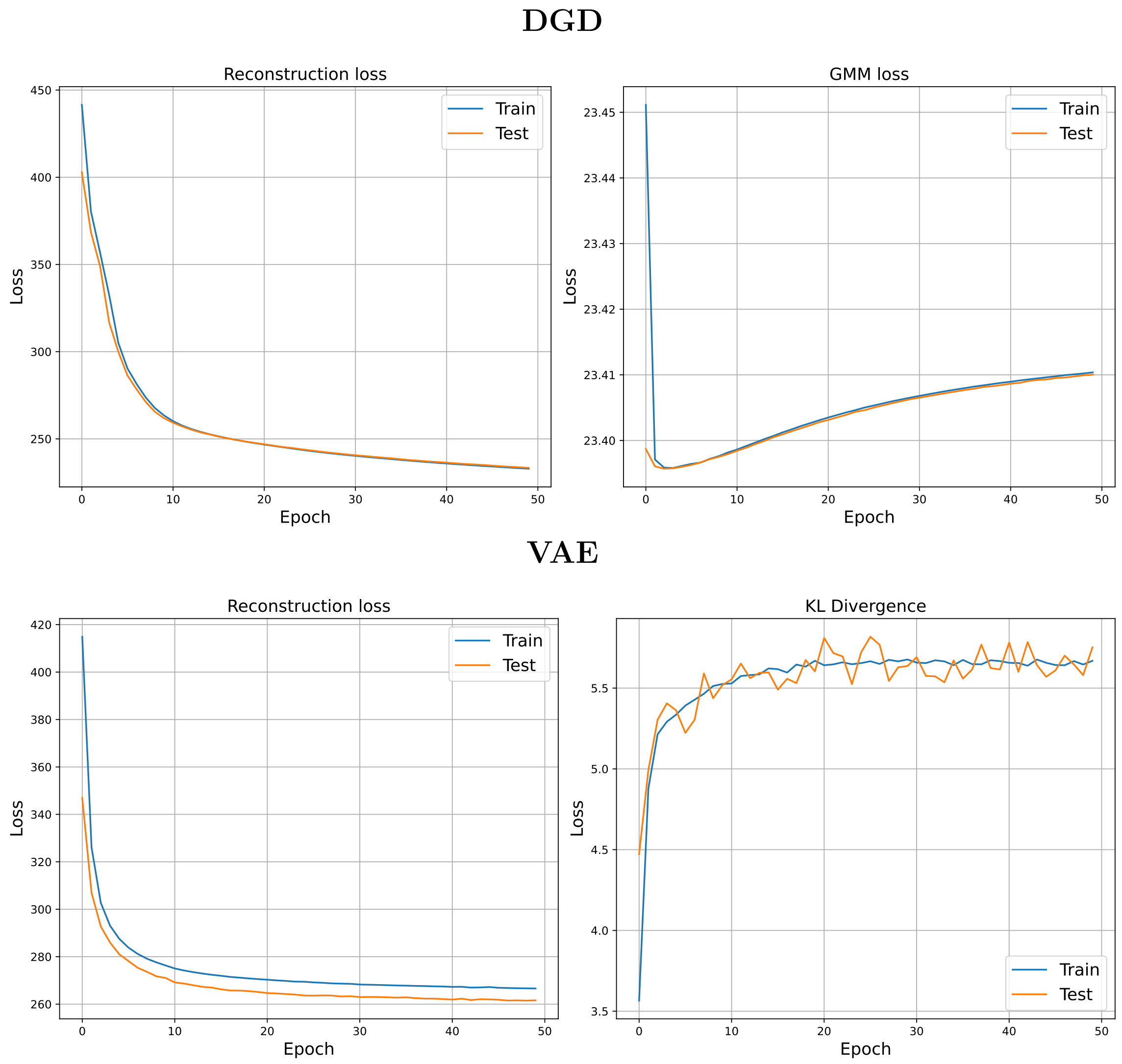


Figure S4: In the top figure, we show the reconstruction loss (BCE) together with the GMM loss for the DGD model, while in the bottom figure, the reconstruction loss (BCE) is displayed alongside the KL divergence loss for the VAE model. For both figures, training losses are represented by blue lines and test losses by orange ones. These curves are referred to the FashionMnist dataset

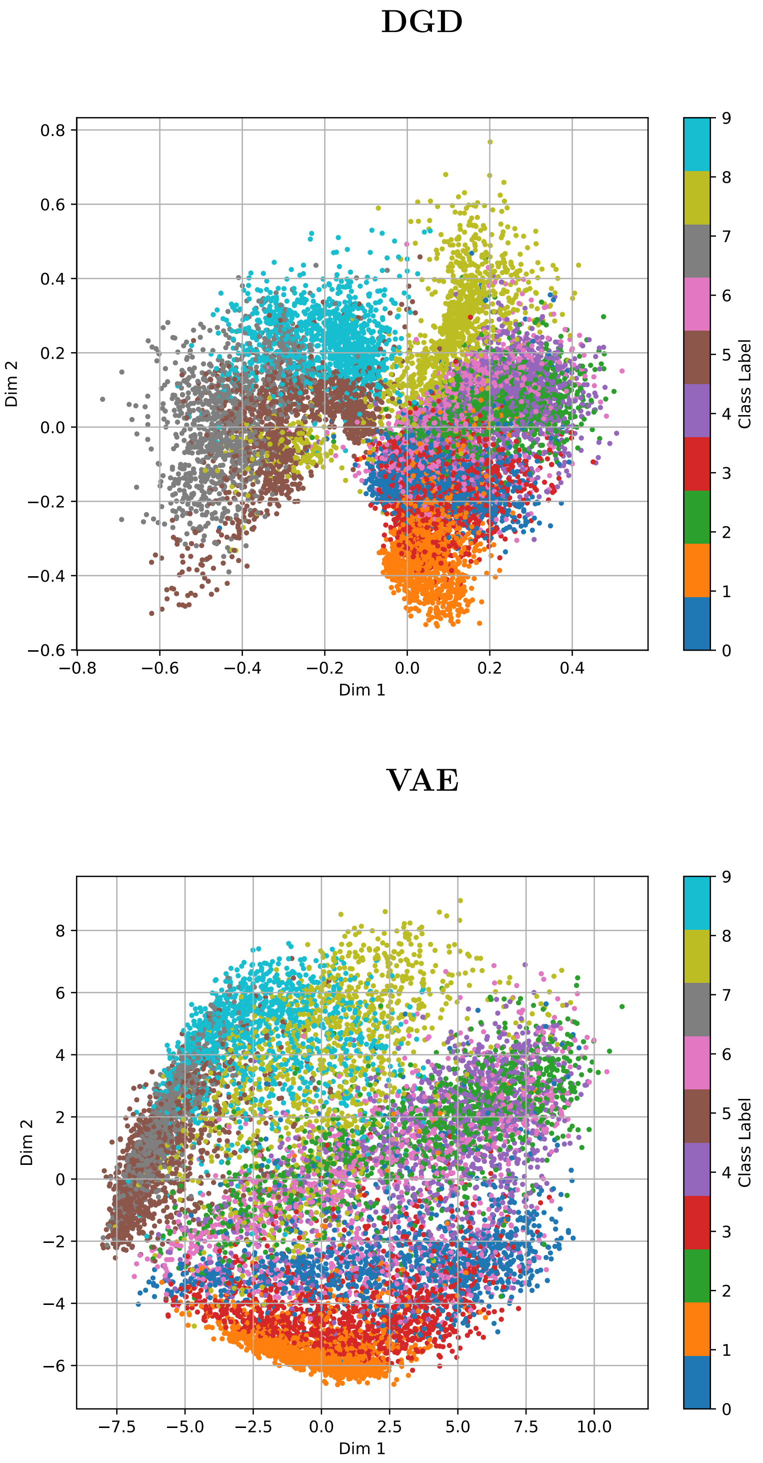


Figure S5: PCA for the latent space of the DGD (top) and VAE (bottom) on the FashionMnist dataset

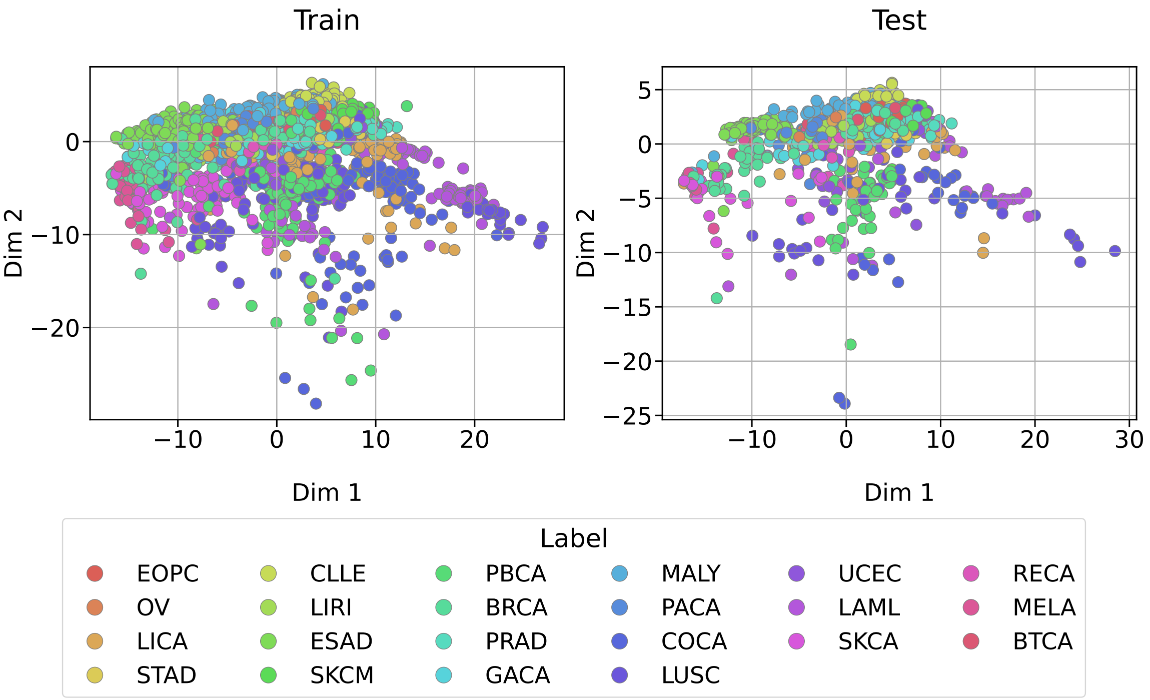


(a)

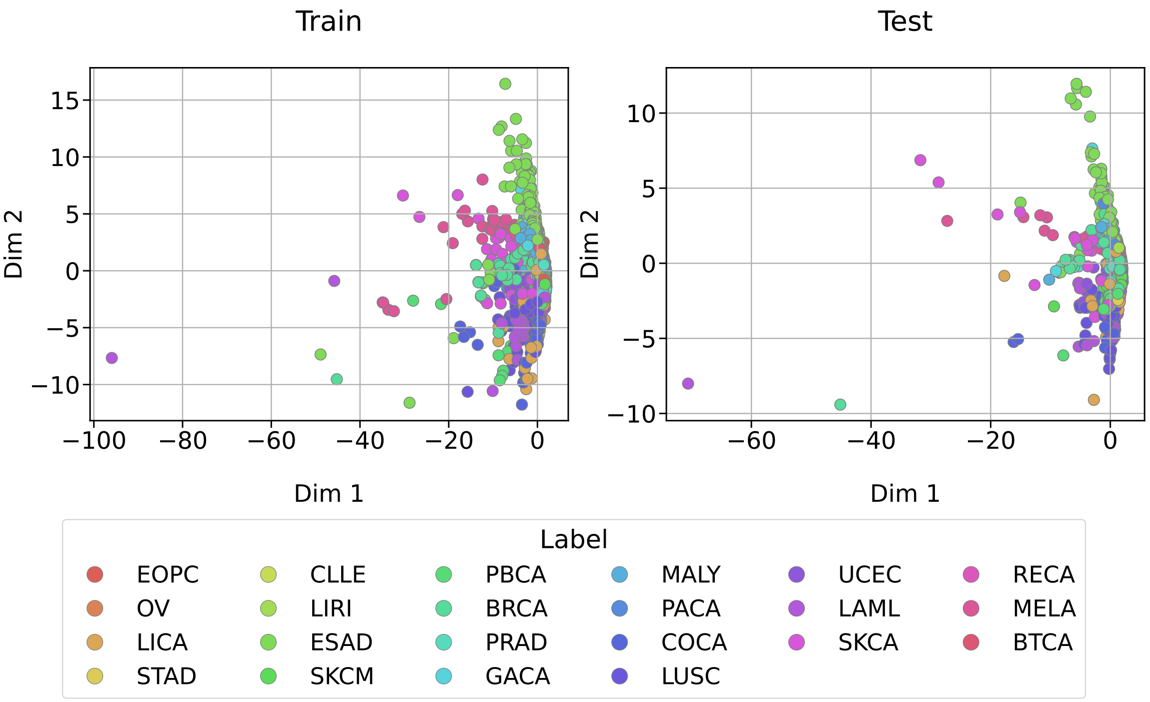


(b)

Figure S6: 2D PCA of the VAE latent representations. (a) PCA projection of the latent space learned by a VAE trained with $\beta = 0$. (b) PCA projection of the latent space learned by a VAE trained using a KL-divergence warm-up strategy.

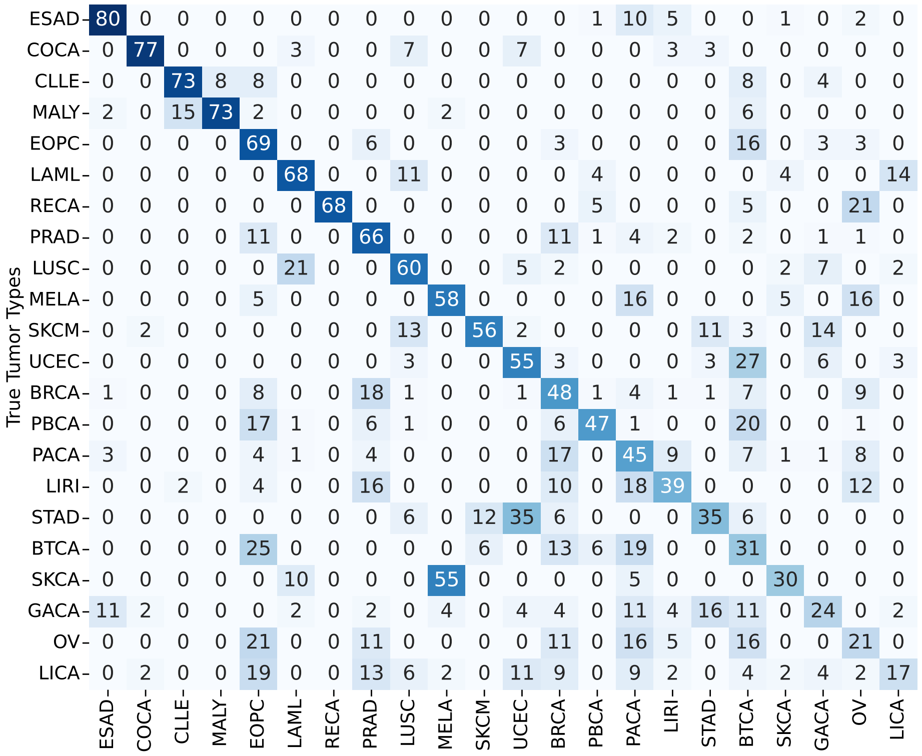


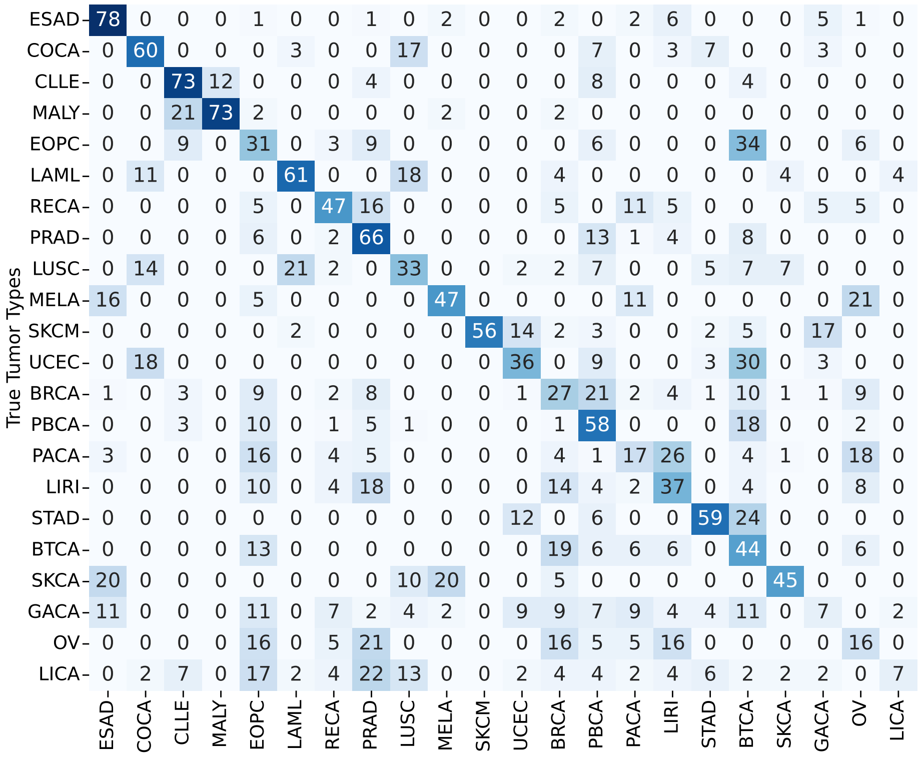


Figure S7: Confusion matrices for the two VAE training strategies. The top panel corresponds to the VAE trained with $\beta = 0$, while the bottom panel corresponds to the VAE trained using a KL-divergence warm-up strategy.
